# Resource competition shapes biological rhythms and promotes temporal niche differentiation in a community simulation

**DOI:** 10.1101/2020.04.22.055160

**Authors:** Vance Difan Gao, Sara Morley-Fletcher, Stefania Maccari, Martha Hotz Vitaterna, Fred W. Turek

## Abstract

Competition for resources often contributes strongly to defining an organism’s ecological niche. Biological rhythms are important adaptations to the temporal dimension of niches, but the role of other organisms in determining such temporal niches have not been much studied, and the role specifically of competition even less so. We investigate how interspecific and intraspecific competition for resources shapes an organism’s activity rhythms. For this, communities of one or two species in an environment with limited resource input were simulated. We demonstrate that when organisms are arrhythmic, one species will always be competitively excluded from the environment, but the existence of activity rhythms allows niche differentiation and indefinite coexistence of the two species. Two species which are initially active at the same phase will differentiate their phase angle of entrainment over time to avoid each other. When only one species is present in an environment, competition within individuals of the species strongly selects for niche expansion through arrhythmicity, but the addition of an interspecific competitor facilitates evolution of increased rhythmic amplitude when combined with additional adaptations for temporal specialization. Finally, if individuals preferentially mate with others who are active at similar times of day, then disruptive selection by intraspecific competition can split one population into two reproductively isolated groups. In summary, these simulations suggest that biological rhythms are an effective method to temporally differentiate ecological niches, and that competition is an important ecological pressure promoting the evolution of rhythms and sleep.

**Author summary:** Why do we sleep? We are interested in the ecological factors which promote the evolution of biological rhythms like the sleep-wake cycle, focusing especially on competition. When animals compete with each other for resources, they often evolve to avoid each other by specializing to use different resources or separating their activity in other ways. To test hypotheses about how competition shapes rest-activity rhythms, we performed computer simulations of a community of animals who move, reproduce, and compete for resources. We show that biological rhythms let two species divide time so that one species is active while its competitor is resting, thus avoiding depleting shared resources. When a species has no competitors in the simulation, competition between members of the same species cause population and individual rhythms to decrease, since resource availability is low when everybody is active at the same time. However, having competitors allows strong rhythms to evolve from originally arrhythmic organisms. Competition can even cause a single population to split into two species which are separated in time. In summary, these results suggest that competition is a strong factor promoting rest-activity rhythms.

## Introduction

An ecological niche is the particular “space” a species occupies in an ecosystem, comprising a diversity of concepts including the physical environment where it lives, the resources it uses, its interactions with other species, and the adaptations it has evolved to exploit the niche. Both theory and evidence indicate that competition is a major factor which determines species’ niches [1,2]. Competitors for a resource generally will adjust to avoid each other, differentiating their niches by adapting to utilize different types of resources, thus avoiding the depletion of shared resources (exploitative competition) [3] as well as costly acts of aggression and territorial exclusion (interference competition) [4].

Time is one dimension which defines a niche. Most species live in environments dominated by a day-night cycle and a seasonal cycle, though other cycles such as tides can also strongly define an ecosystem [5]. A temporal niche is defined by both abiotic elements such as light intensity and temperature [6,7], as well the as by predictable temporal patterns of other organisms [8]. Adaptions for a specific temporal niche include for example specializations in vision, coat color, temperature regulation, and importantly, biological rhythms, i.e. physiological and behavioral traits which cycle regularly according to an internal biological clock. In this article, we consider activity rhythms through the lens of ecological niche theory, viewing them as a way partition niches among competitors and examining how competition shapes the expression of these biological clocks. Time can be thus viewed as a sort of resource or space which is partitioned by organisms in response to competition [9], both interspecific (between different species) and intraspecific (between members of the same species) [10]. While some work has examined predation’s effect on animals’ activity rhythms [11–13], the role of competition has been less examined.

A variety of field studies have observed evidence of temporal partitioning between putative competitors. For example, a community of three sympatric species of insectivorous bats were observed to have activity profiles which are displaced from each other, suggesting that they avoid each other [14]. While Argentinian pampas foxes are normally nocturnal, they show a modified half-nocturnal half-diurnal profile in a region where they overlap with crab-eating foxes [15]. Other examples of apparent temporal partitioning have been seen at watering holes [16,17], in desert gerbils [18], carabid beetles [19], and large African carnivores [20]. Stronger evidence comes from semi-natural manipulation experiments. Common spiny mice are nocturnal and golden spiny mice are diurnal in areas where the two species coexist, but when the common spiny mice are experimentally removed, the originally diurnal golden spiny mice shift into nocturnality, suggesting that this species was pushed into diurnality by the competing species [9]. It should be recognized that these examples all occur at the behavioral and ecological time scale, i.e. they are “temporary” displacements from the species’ genetically determined preferred activity time. Although certain pairs of taxa are suggestive of evolutionary-level temporal partitioning—nocturnal owls and diurnal hawks, for instance, or the hypothesized nocturnal origin of mammals [21]—the role of competition in causing these evolutionary divergences is near impossible to establish conclusively. Nonetheless, it is probable that temporal shifts at the behavioral/ecological level cause selection for alleles suited for the new time period, eventually translating into long-term changes at the genetic/evolutionary level.

In this work, we explore ways which competition in ecological niche theory can be applied to biological rhythms, using simulation of a community of two species who compete for limited resources in an environment. We first demonstrate that circadian rhythms are a form of niche differentiation which allows two species to stably coexist without competitive exclusion, and that two rhythmic species will quickly differentiate their circadian phases to avoid competition. We show that arrhythmicity is preferred when there is no interspecific competitor present, but the presence of interspecific competition greatly facilitates the development of strong activity rhythms. Finally, we show that if organisms have assortative mating based on circadian timing, then intraspecific competition can drive the division of one population into two mating groups, contributing to speciation.

## Results

Briefly, we simulated a community of two species in an environment using Python. Each individual organism has a circadian activity rhythm which is modeled as a cosine wave with two parameters: amplitude ranging from 0 to 1, and phase angle ranging from 0° to 360° (Fig 1). Resource input into the environment occurs at a constant rate. If an individual encounters a resource, it consumes that resource and gains energy, which is in turn used for movement and basal metabolism. Death occurs if an individual’s energy drops to zero or if the individual reaches an age limit. Organisms reproduce sexually, and the probability of an individual reproducing is proportional to its energy. During procreation, an offspring’s amplitude and phase are the means of the parents’ traits. When testing the evolution of a trait, the offspring’s trait is the parents’ mean plus or minus a small random variance, which naturally results in natural selection and evolution.

**Fig 1.**
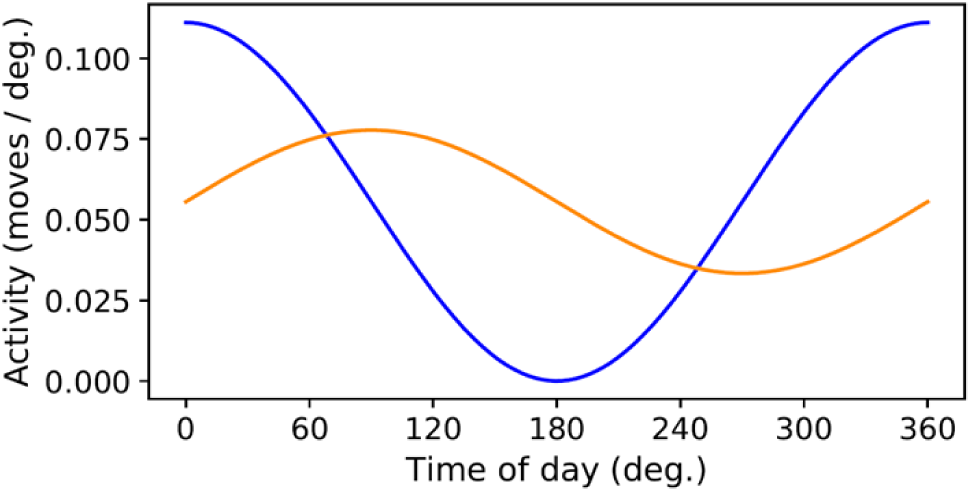
Example activity rhythm profiles. Activity profiles are modeled as cosine waves with properties amplitude and phase angle. Two example profiles are shown. In blue, the activity rhythm has amplitude = 1 and phase angle = 0°. In orange, amplitude = 0.4 and phase angle = 90°. Regardless of changes in circadian parameters, the number of moves per day is fixed at 20 (equivalent to the area under each curve).

### Rhythmic activity prevents competitive exclusion

The competitive exclusion principle asserts that two species cannot occupy the same ecological niche indefinitely; if two species exist in exactly the same niche, then one species will eventually outcompete the other and become the sole occupier of the niche. We modeled this scenario by simulating two species with no circadian rhythms, i.e. with amplitude = 0. The two species are constantly active at a moderate level and thus have no differentiation of temporal niches. In this situation, one species invariably dies out (Fig 2A-B). If both species are rhythmic (amplitude = 1) and are active at the same time of day (same phase angle), there is again no niche differentiation, and one species will eventually die out (Fig 2C-D). However, if both species are rhythmic but are active during opposite times of day, then both species are likely to continue existing (Fig 2E-F). In 2 of the 40 trials of this scenario, one species died out after some time, demonstrating that random fluctuations can still cause a species to disappear. In the remaining 38 trials, both species survived at the end of 2000 days.

**Fig 2.**
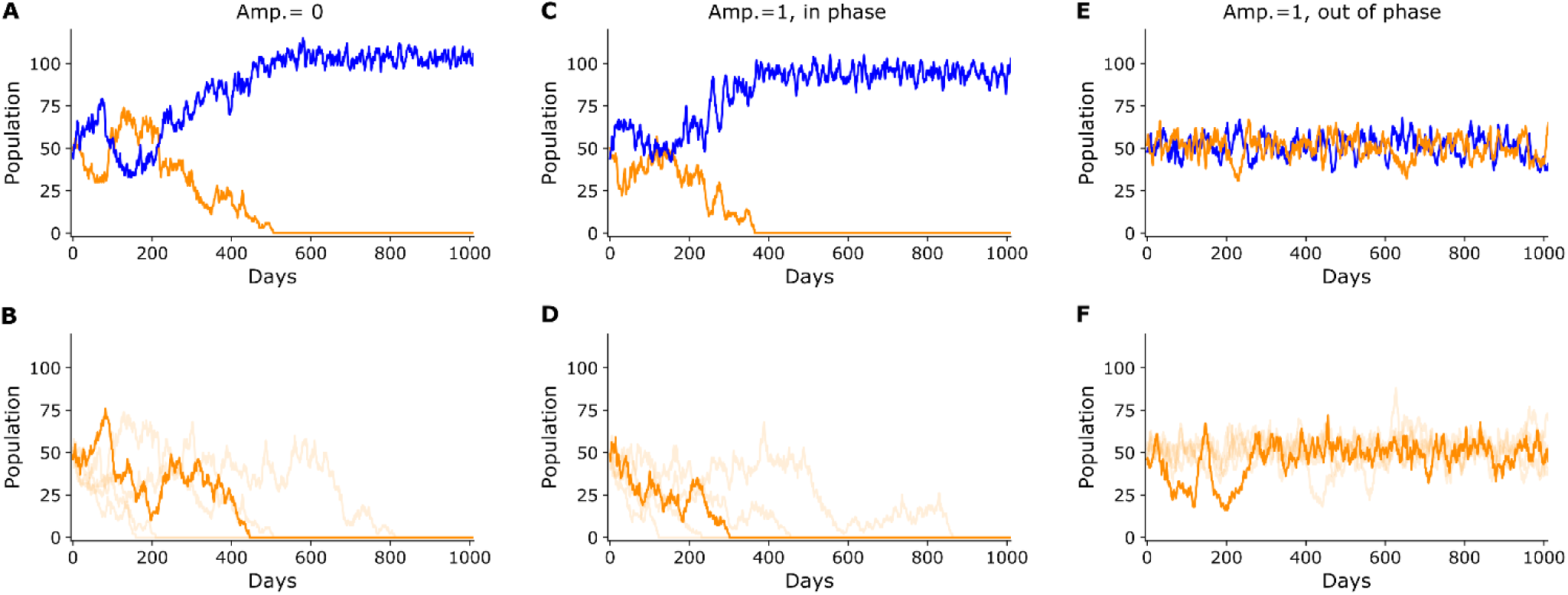
Circadian rhythms prevent competitive exclusion. (A) When two species with no activity rhythms (amp.=0) are placed in the environment, one species will always die out. An example of an individual trial is displayed. (B) Summary of 40 trials. Since the two species’ population levels tend to be symmetric, only the species which is excluded is plotted for clarity. The trials were sorted by the day that competitive exclusion is reached, and the median (20^th^) trial is represented by the dark line, while the 4^th^, 12^th^, 28^th^, and 36^th^ trials are represented by faint lines. (C) When two species have circadian rhythms (amp.=1) and are active at the same phase, one species will always die out, as before. An example individual trial is displayed. (D) Summary of 40 trials. (E) When two species have circadian rhythms are active 180° out of phase, the two species can coexist indefinitely. An example individual trial is displayed. (F) Summary of multiple trials. Since no species died out in this scenario, five population profiles were selected at random to be plotted.

### Character displacement of phase-angle

Having showed that being active at opposite phase angles is favorable for the coexistence of two species, we ask whether species can evolve over time to reach such a state of temporal separation. When two similar species occupying similar niches are found in the same geographical area, the differences between them tend to become accentuated in order to distinguish their niches, in a phenomenon known as character displacement [22]. We tested whether two circadian-rhythmic species would display character displacement on a temporal axis. In this scenario, all individuals started at approximately at the same phase angle at the beginning of the simulation, and the offspring’s phase angles were allowed to vary and thus evolve, while amplitudes were fixed at 1.

The two species’ phase angles of activity did indeed diverge over time in the simulation (Fig 3A). After starting at 0° separation, the phase-angle difference between the two species grew and eventually reached the maximum of 180° and remained so thereafter (Fig 3B). The source of selective pressure is the difference in resource availability between peak and peak-adjacent times (Fig 3C). During times of peak activity, available resources are quickly consumed and resource density is thus very low, while in times adjacent to the peak, less resources are consumed, and resource density is higher. The result of this time-resource gradient is that individuals whose phase angles are at the edges of the population distribution have access to a more resource-dense environment and thus gain more energy (Fig 3D). Since stored energy is proportional to reproduction probability, the individuals at distribution’s edges reproduce more, and the species’ mean phase angle is shifted.

**Fig 3.**
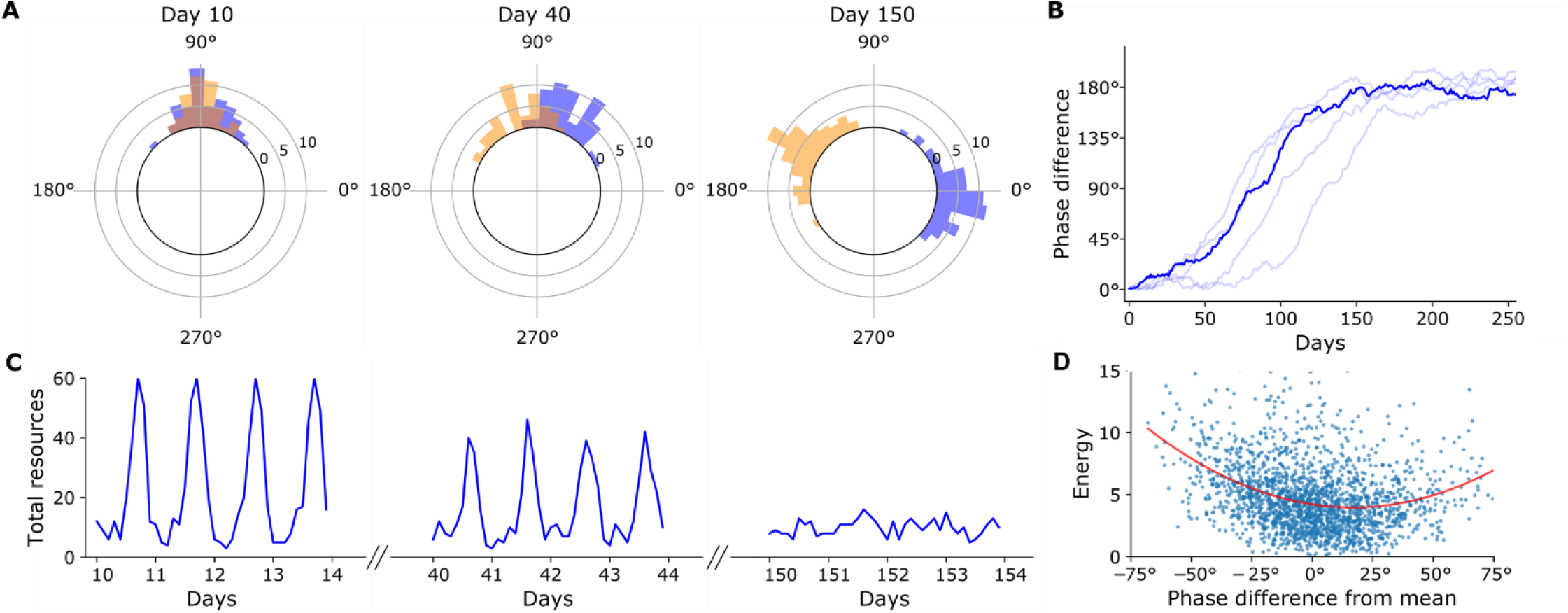
Character displacement of circadian phase. (A) Circular histogram displaying the separation of phase angles of activity in two species over time in one trial. The difference in the two species’ phase angles of entrainment quickly diverges to 180°. (B) Summary of 40 trials. Trials were sorted by days until a phase difference of 150° was reached. The 20^th^ trial is plotted in a dark line, while the 4^th,^ 12^th^, 28^th^, and 36^th^ trials are plotted in faint lines. (C) When the two species are at similar phase angles of entrainment, there is a large difference in total resources in the environment during different times of day. When the two species are separated in phase, there is no longer a predictable cycle in resource availability. (D) Scatterplot of organisms’ stored energy vs their phase angle of entrainment relative to the population mean. The parabolic best fit line is shown in red. Individuals at the edges of the population phase distribution acquire more energy and consequently are more likely to mate.

### Intraspecific competition causes temporal niche expansion by arrhythmicity

Previous work has shown that when interspecific competition is absent, the population’s niche width tends to expand, as competition within the species pushes individuals to exploit a greater range of resources [23]. We tested if this pattern would be true in this model. Indeed, when rhythmic amplitude was allowed to vary, the amplitdue steadily decreased to near-zero levels, representing the transition from a temporal specialist to a generalist strategy. To clarify, a broad circadian niche does not necessarily imply the absence of sleep or sleep-like rest; these organisms could follow a cathemeral pattern in which sleep is scattered throughout the day.

However, other factors may prevent dampening of rhythms. To adapt to a temporal niche, organisms do not only develop activity rhythms but additional morphological and physiological specializations as well, such as in vision, temperature regulation, and non-locomotor circadian rhythms [9,24]. These adaptations allow an animal to acquire more resources per unit energy expenditure during their specialized “on” phase, but often at the cost of being less efficient during their “off” phase. A temporally specialized organism would thus incur costs when adopting a generalist activity rhythm. To model these temporal adaptations, we added a specialization term to the simulation. The degree of specialization over the day follows a cosine wave such that at specialization = 0.1, an animal acquires an additional 10% energy from consuming a resource during the peak and 10% less energy during the trough. For simplicity, we assume that the phase angle of the specialization wave matches exactly that of the cosine wave of circadian activity, as opposed to modeling them as separate phenotypes, which would naturally converge to match phase angles anyways so that organisms are most specialized when they are most active.

We found that specialization must be quite high to overcome competition’s negative selection for amplitude (Fig 4). At low levels of specialization, the equilibrium amplitude is positively correlated with the degree of specialization. Only when specialization is at least 0.75 does the resulting amplitude approach 1 (Fig 4C). This is rather intense specialization, indicating that resource acquisition is 7 times more efficient during the peak than during the trough. This suggests that, in an environment where resource availability is roughly constant throughout the day, intraspecific competition is a strong factor selecting for arrhythmicity.

**Fig 4.**
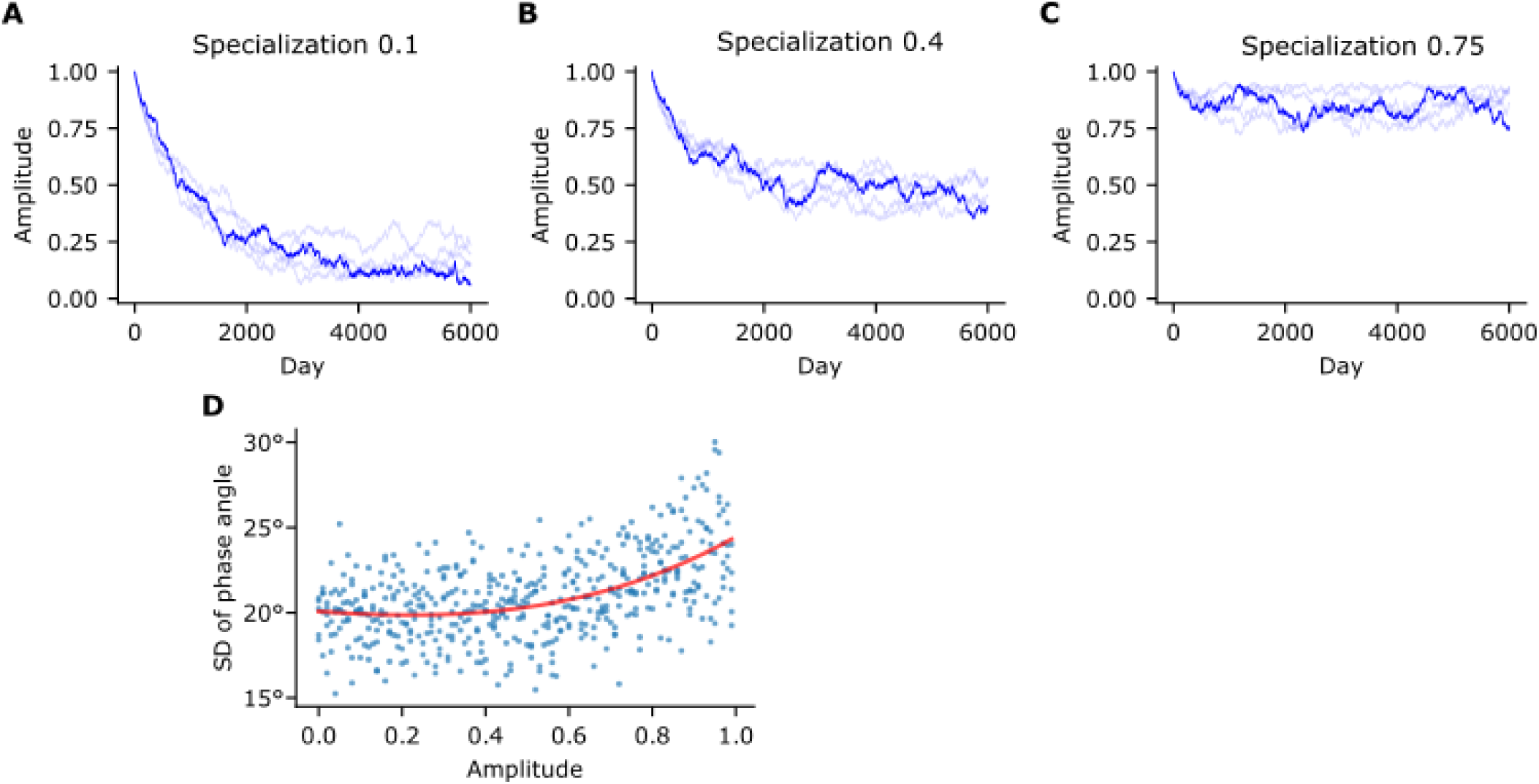
Intraspecific competition selects for arrhythmicity. (A) Even with a positive specialization value, amplitude is driven towards low levels when there is only one species. (B) Amplitude settles on an intermediate value when specialization = 0.4, demonstrating a positive correlation between specialization and resulting amplitude. (C) In the single-species scenario, specialization must be set quite high at 0.75 for amplitude to reach near 1. (D) Standard deviation of phase angle as a function of amplitude. This illustrates an inverse relationship between expanded interindividual niche width (high phase angle variance) and expanded intraindividual niche width (low amplitude).

When a population is freed from competition, the following niche expansion could occur through increasing the niche width within individuals, or through increasing the variance in between individuals who are still relatively specialized. Whereas low rhythmic amplitude represents within-individual niche expansion, we also observed between-individual niche expansion, represented the greater range of phase angles when amplitude is high (Fig 4D). When amplitude is low, there is no selective pressure to promote interindividual variance, so sexual reproduction pulls the population towards the mean.

### Intraspecific competition allows rhythmicity to develop

While intraspecific competition acts to prevent rhythmicity, we asked if a competitor species would cause rhythmicity to develop. When specialization = 0, organisms did not show a trend towards either high or low rhythmicity (Fig 5A). Since the two species’ activity rhythms were in approximately opposite phase and similar amplitude, the summed activity of all individuals in the environment was approximately constant throughout the day, which in turn caused resource availability to be approximately constant. This situation is much like the final condition of Fig 3A and Fig 3C. With no time-resource graident, there is no evolutionary pressure for rhythmicity to evolve.

**Fig 5.**
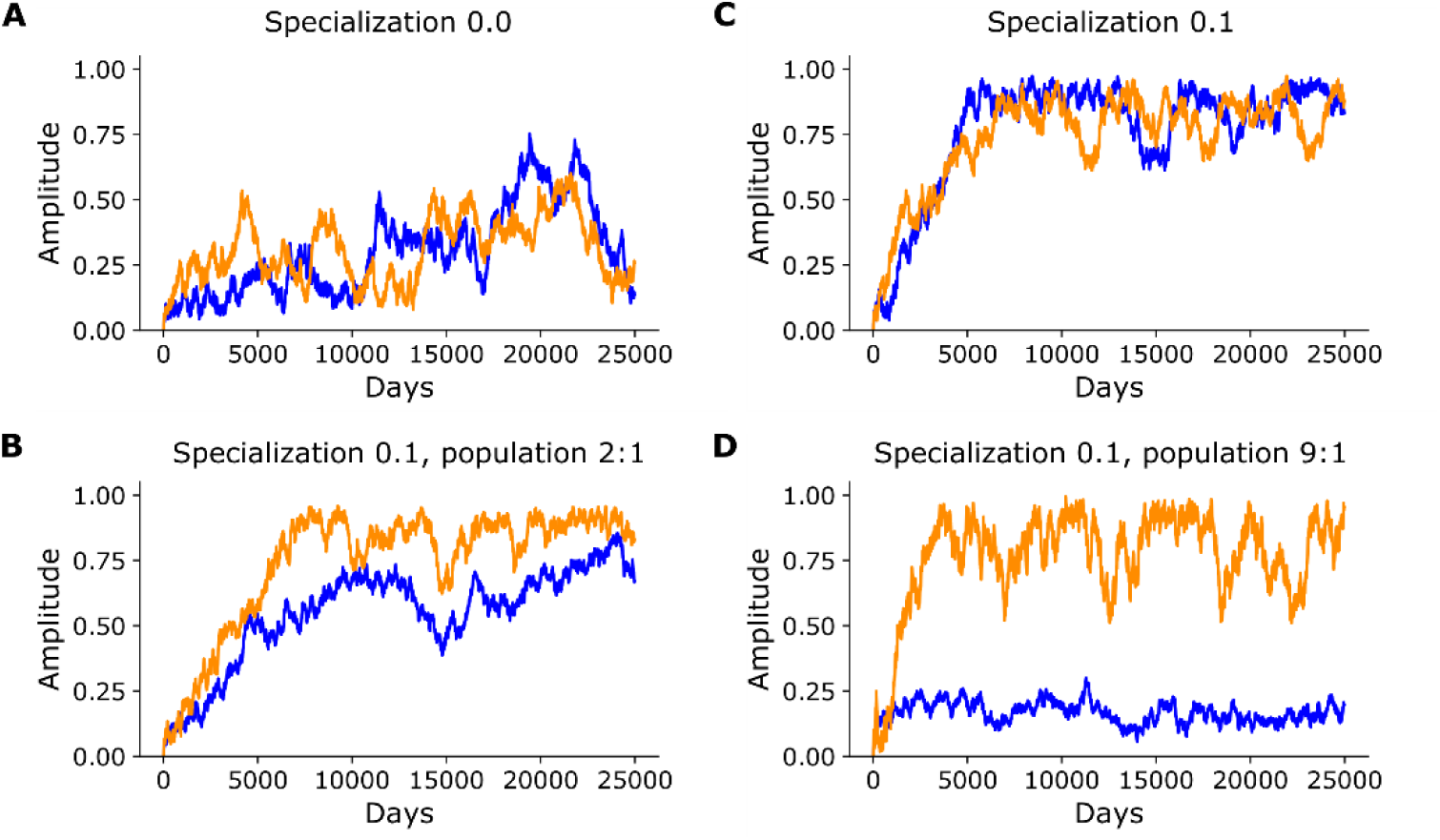
Interspecific competition with specialization allows development of biological rhythms. (A) With two species, amplitude drifts randomly without pressure towards either 0 or 1. (B) When a specialization value is added to the model, amplitudes rise steadily over time. (C) The final equilibrium amplitudes depend on relative population numbers. When the blue species is twice as numerous as the orange, its final amplitude remains lower than the orange’s. (D) When the blue species is nine times as numerous as the orange, blue’s amplitude remains low while orange’s is high.

While interspecific competition alone does not positively select for amplitude, it also effectively neutralizes intraspecific competition’s negative selection for amplitude. In this state, if the organisms have even a small degree of specialization, it can drive amplitude to increase steadily over time in both species (Fig 5B), with higher specialization values causing amplitude to increase more quickly. Whereas a specialization value of 0.1 causes amplitude rise to 1 when there are two species, amplitude barely rises when there is only one species (Fig 4A), and a specialization value as extreme as 0.75 is needed to raise amplitude to the levels seen in the two species scenario (Fig 4C).

The amplitudes of the two species tended to be correlated (Fig 5A-B). The correlation of the two species’ amplitudes can be explained by the fact that a chance increase in amplitude in one species causes a resource density difference in time, which encourages the other species to increase their amplitude due to interspecific competition and the first species to decrease their amplitude due to intraspecific competition.

The relative strength of interspecific and intraspecific competition influences the equilibrium amplitude of each species. To simulate asymmetric populations, we defined a target population ratio, and the populations were stabilized around this ratio by automatically adjusting their reproduction rate. When species A is set to be two times as numerous as species B, species B develops high rhythmicity while species A’s amplitude hovers around 0.7 (Fig 5C). If species A is set to be nine times as numerous as species B, species A’s amplitude remains only at 0.2, demonstrating that intraspecific rather than interspecific competition is the much stronger force in determining its equilibrium amplitude. This also demonstrates that a dominant species which has even mild rhythms can strongly determine the rhythms of minor competitors which have smaller populations. In total, these results suggest that the presence of competitors is a strong factor promoting the expression of biological rhythms.

### Population splitting by differentiation of phase

The single-species scenario illustrated that intraspecific competition causes expansions in both within-individual and between-individual niche widths. Could an increase in between-individual variation cause the population to split into two distinct groups? This would in effect cause speciation, or at least contribute strongly to the ecological divergence and reproductive isolation which are precursors to full speciation. To test this, assortative mating was introduced to the model. Individuals only mate with a partner whose phase differs by less than 9 hours (135°), and the probability of choosing a given partner increases linearly as the phase difference decreases to zero. Amplitudes were fixed at 1 for this test.

We found that, with the addition of assortative mating, the population did indeed segregate into two groups (Fig 6A). We defined a cross-group mating value (CGM) to quantify reproductive isolation, where low CGM indicates high reproductive isolation. The CGM tended to stay constant for some time and then drop suddenly (Fig 6B). This suggests that there is a threshold effect, such that once the conditions occur for a secondary population to escape “gene flow” with the main population, full reproductive isolation quickly follows. To separate from the main population, there must be enough individuals at the far edges of the phase distribution to sustain a “breeding population”, such that a tipping point is reached where individuals in the incipient secondary population become more likely to breed within its own population rather than the main population. Once this tipping point is reached, the secondary population quickly splits off from the main group. Given the random nature of mate selection and progeny’s phase variance in our model, the conditions where this happens occurs stochastically, explaining the large variance in speciation time in the simulations.

**Fig 6.**
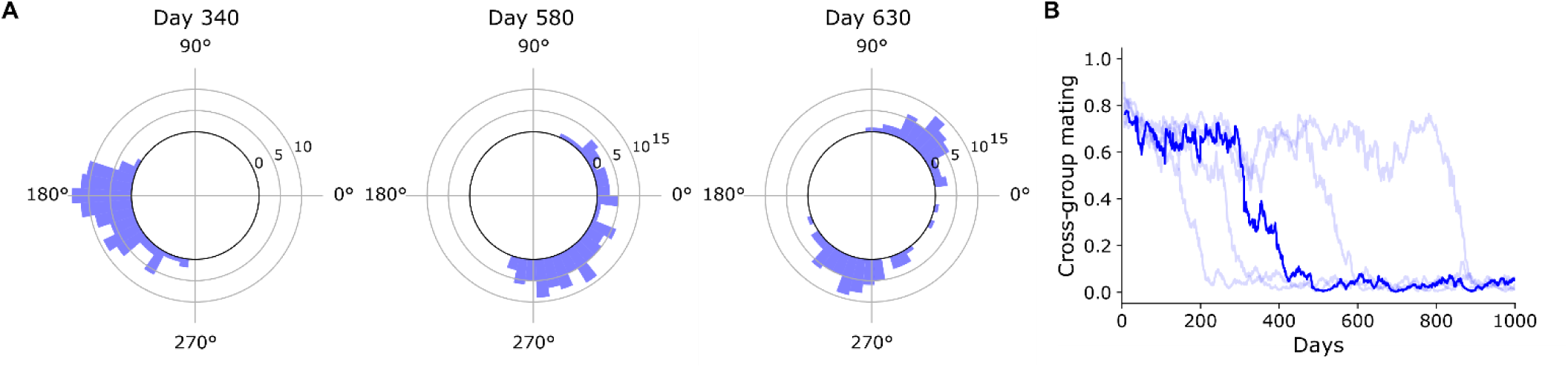
Population splitting by phase. (A) Circular histograms showing distributions of phase angles of entrainment in one trial. On day 340, The population was a single mating group (cross-group mating = 0.67). On day 580, the population is in the process of splitting into two mating groups (CGM = 0.38). By day 630, there are two populations which are almost completely reproductively isolated (CGM = 0.05). (B) Summary of 40 trials. Trials were sorted by days to reach CGM < 0.1; the 20^th^ trial is plotted in dark blue, while the 4^th^, 12^th^, 28^th^, and 36^th^ trials are plotted in light blue.

Population splitting was not guaranteed for all parameter settings. Conditions necessary to achieve speciation in this model are somewhat constrained, which is unsurprising given the high barriers to sympatric speciation in nature [25]. The time-resource gradient must be quite strong to reach the necessary levels of density-dependent disruptive selection, and thus population splitting will not occur for low resource regeneration rates or low population levels. A smaller mating window of course also promotes reproductive isolation. Higher variance in offsprings’ phases also promotes reproductive isolation by increasing the chance of a breakaway group to escape “gene flow” with the rest of the population. A long lifespan also favors speciation, since individuals at the edges of the distribution then have more time to build up an energy surplus.

### Population splitting vs. individual expansion

Intraindividual niche expansion will tend to oppose speciation, since a decrease of rhythmic amplitude would weaken the time-resource gradient and broaden mating windows. Conversely, speciation into two populations occupying opposite niches will remove selective pressure for broad individual niches since the time-resource gradient would be erased (analogous to the last case in Fig 3C). Faced with intraspecific competition, which of these two mutually exclusive paths will a population take? Results suggest that it depends heavily on the relative flexibility of the amplitude or phase traits. We set standard deviations of phase angle and amplitude to various test values, with a higher standard deviation resulting in quicker evolution of that trait. When phase standard deviation was high and amplitude standard deviation was low, populations were more likely to speciate; conversely, when phase standard deviation was low and amplitude standard deviation was high, populations tended to become arrhythmic (Table 1).

**Table 1.** Number of trials achieving speciation depend on the variances of amplitude versus phase angle.

|  |  | Amplitude standard deviation |  |  |  |
| --- | --- | --- | --- | --- | --- |
|  |  | 0.02 | 0.03 | 0.04 | 0.05 |
| Phase standard<br>deviation | 0.04 | 10/20 | 0/20 | 0/20 | 0/20 |
|  | 0.045 | 20/20 | 14/20 | 9/20 | 5/20 |
|  | 0.05 | 20/20 | 20/20 | 17/20 | 12/20 |
|  | 0.055 | 20/20 | 20/20 | 20/20 | 17/20 |

## Discussion

This is the first modeling study to examine the ecological pressures which favor the evolution of biological rhythms. While other modeling studies have examined the “internal” forces which make rhythms and sleep physiologically efficient [26–28], none have looked at the “external” forces of the environment.

### Timescales of analysis

Though we framed our model in terms of organisms evolving genotypically over time, niche partitioning can equally occur at the timescale of individual behavior or ecological community [29], and the conclusions of this study can easily be generalized to sub-evolutionary timescales and mechanisms. Many animals are known to switch their temporal niches depending on various environmental factors [24]; for example, the aforementioned studies on minks, spiny mice [30], and foxes [15] demonstrate temporal partitioning at the sub-evolutionary level. Indeed, every clear observation of competition’s effect on temporal niches has occurred at the behavioral/ecological level, and inferring the role of competition in a species’ evolution is difficult if not impossible, since those historical interspecies interactions are unobservable. However, if temporal displacement continues for an extended time, it seems likely that organisms would eventually evolve to match their internal clock with their expressed behavior. Research on the health consequences of misalignment between the circadian clock and activity rhythms [7,31,32] suggest clear selective mechanisms by which behavioral displacements can translate into genetic/evolutionary changes.

The simulation was framed in terms of circadian rhythms, but the results could equally apply to seasonal rhythms. A major difference between the daily and seasonal timescales is that individuals can adjust their daily behavior to a new time through non-genetic mechanisms, whereas this is probably less common at longer timescales, though it certainly depends heavily on the particular mechanisms governing these rhythms.

### Competitive exclusion and character displacement

In our model, we demonstrated that competitive exclusion is unavoidable if two species share an ecological niche. Two species which are temporally separated are much more likely to coexist for extended periods of time. Additionally, we show that two circadian-rhythmic species which initially are active at the same phase evolve to be active at opposite times of day in order to avoid competition for resources. The most likely direct natural analog of this scenario is the introduction of a new species to a region already occupied by another similar species. This could happen, for example, through the breakdown of geographical barriers, chance migration to a new area [33], introduction by humans, or climate change-induced range shifts [34]. The invasive American mink in the United Kingdom offers a good real-life example: originally nocturnal when first introduced, the mink largely switched to diurnality when the native otters and polecats recovered their population levels [35]. The previously mentioned studies in Argentinian foxes [15] and desert spiny mice [9] also demonstrate niche switching.

### Intraspecific competition and niche expansion

Intraspecific competition is hypothesized to increase the niche width in a population, especially in environments where a population is released from competition from other species, which could occur after migration or introduction to a new region, for instance [36,37]. The single-species simulations demonstrate this phenomenon for temporal niches.

Niche expansion can happen either by increasing the variance of resource type usage between individuals [38,39] or by increasing the range of resource usage of each individual [40]. Interindividual expansion appears to be the more common pattern in nature in general [41], especially if intraindividual generalization has associated costs [23]. Our single-species simulation displayed both interindividual variation, visible as a broadened phase-angle distribution, and intraindividual generalization, visible as flattened amplitude. The model’s design does not allow for very extreme interindividual expansion, since activity timing was determined strictly by genetics and not by individuals’ behavioral adjustments, and sexual reproduction tended to pull genotypes towards the population mean. Programming a non-genetic behavioral component may be interesting for future studies.

A few studies in fish have observed offset patterns of diel feeding activity between individuals, with larger dominant individuals feed at preferred times while subordinates being pushed to other times [13,42]. This suggests that preexisting heterogeneities in the population such as social dominance could predispose niche expansion to occur between rather than within individuals. We are aware of only one study to observe temporal niche expansion in the wild; American minks which were introduced to a remote island with no competitors were observed to have expanded temporal niches compared to their mainland counterparts [43]; it is unknown whether this was due more inter- or intraindividual expansion. Intraspecific competition is likely to also play a role in instances of niche expansion after release from predation [11]. Predator-free space may be seen as a type of resource, which becomes more abundant after removal of the predator. Though never specifically examined, it is reasonable to suspect that intraspecific competition could then drive the subsequent niche expansion observed after predator release.

Arrhythmic/cathemeral activity patterns appear to be somewhat common [44,45]. However, besides the aforementioned observation of invasive minks [43], there have not been any other experiments or observations to directly test competition’s role in causing temporal niche expansion. We suggest that this phenomenon is not uncommon and may explain the behavior and evolution of a number of these arrhythmic animals.

### Competition facilitates emergence of rhythms

The presence of an interspecific competitor erases the advantage that arrhythmicity in had in a single-species scenario. A small amount of time-of-day specialization (theoretically, any amount of specialization) can then cause organisms to evolve high amplitudes. We only modeled exploitative competition here, where fitness differences were only driven by differential access to resources. If interference competition were also present, for example if a cost was incurred every time an individual encountered an allospecific, then development of rhythmic amplitude would most likely occur even more quickly.

As previously mentioned, the comparison between invasive minks on an island and mainland minks is an instance of apparent competition-mediated niche expansion/contraction [43], but we are not aware of any studies which has observed competition inducing increased amplitude. Again, we believe this phenomenon does exist broadly, but has simply not been observed yet. Another potential real-life analog is if two arrhythmic species existing in a constant environment—cave-dwelling fishes for example [46]—are suddenly exposed to a cycling environment and are thus pushed to temporally differentiate.

### Speciation

When individuals were limited to mating with other individuals who were active at similar times of day, intraspecific competition could cause a single population to separate into two groups which were largely reproductively isolated, in effect causing speciation. This is specifically a case of *sympatric* speciation, in which new species arise without geographic separation. Speciation required high levels of density-dependent selection (i.e. a strong time-resource gradient) to occur. This is not surprising given that sympatric speciation in general is believed to be quite difficult to achieve in nature, such that its existence at all was questioned before recent decades [25]. A theoretical problem against sympatric speciation is that the phenotype which improves fitness usually has no effect on mate choice, and thus even when disruptive selection exists, there is no mechanism for it to translate to mating preferences. However, speciation by time (allochronic speciation) has an advantage here because it is a “magic” trait [47], i.e. it has fitness effects which are acted on by natural selection, *and* it is also directly responsible for assortative mating.

A recent review [48] finds good evidence for dozens of allochronic speciation events, with most cases occurring at the seasonal rather than the circadian timespan, but although intraspecific competition is known to play a role in promoting speciation in general [49,50], it is unclear how important intraspecific competition was in these specific cases of allochronic speciation. In a study of Madeiran storm petrels, the authors hypothesize that competition for food and nest space may have led to allochronic separation into hot- and cool-season breeding populations [51]. Besides competition, allochronic speciation can occur by other reasons too, such as temporal isolation [52], environmental changes [53], or a founder effect [54]; temporal separation can also be secondary to another adaptation [55] or serve to reinforce reproductive isolation. Intraspecific competition can act alongside these other mechanisms to produce allochronic speciation.

### Sleep

The question of why sleep exists is one of the big questions of biology. Traditionally framed something of a paradox that an animal would abandon motor and sensory activity for several hours per day, it is perhaps not so mysterious after all when considering that temporal niches exist. External pressures, whether they be competition or other environmental forces, push animals to specialize for a time of day. Internal pressures, such as energy conservation [56], the need to perform restorative functions, and the separation or coordination of clashing or synergistic physiological processes [27,28,32], also encourage the evolution of sleep (or at least a rest period). Especially with external and internal pressures combined, sleep/rest appears to be an unsurprising evolutionary consequence. Stated colloquially, “if I can’t be active right now, then I might as well rest. If rest is efficient, then I might as well do it when I can’t be active.”

As to why a defined sleep state exists rather than simply circadian-mediated inactivity, we propose that true sleep can be viewed as an adaptation to delay rest. Urgent circumstances sometimes interrupt an animal’s regularly scheduled rest, and thus the defining feature of true sleep, sleep homeostasis (“sleepiness”), encourages animals to later pay back the lost rest. Another feature of true sleep is that it is a relatively distinct mode with a well-defined sleep-wake boundary. In unpredictable environments, such a clear distinction allows animals to maximize rest when the opportunity is available, rather than a slow transition between rest and sleep.

### Methodology and future directions

We intentionally created a model without distinct day and night phases, so that it would be more generalizable to a variety of hypothetical conditions. Nevertheless, for most organisms, the changes between day and night represent a major division in the environment [44], and thus alternative models may explicitly include this. Activity patterns other than cosine waves, or testing the variable of daily total activity, could also be interesting.

We decided not to use a genetic model of circadian rhythms, opting instead to model only phenotypes with a constant distribution. We do not believe this is a weakness because it is more or less equivalent to a situation of polygenic inheritance with many genes. However, a genetic model may have some advantages such as allowing a clearer look at population genetics. Additional models have been developed to explain the theoretical benefits of sleep and circadian rhythms from the point of view of internal physiological organization [27,28], and it could be interesting to integrate models of these “internal” benefits of rhythms with the “external” ecological benefits modeled here.

Many questions regarding this topic remain open for investigation. Among the many possible axes by which competing species can differentiate, what promotes partitioning of time over partitioning of other resources? How does competition interact with other forces, like predation, to influence rhythms? To what extent does competition shape activity rhythms in nature? In addition to additional field studies, well-designed experiments with lab rodents could even help us understand the circumstances in which activity rhythms are modified by competition.

## Methods

### Environment and organisms

Using Python, we simulated communities of organisms existing in an environment consisting of 60 spaces, where each space may contain a resource, contain an individual organism, contain multiple individuals, or be empty. An individual moves by jumping to another random space, and if it moves to a space which contains a resource, the resource is consumed.

600 resources per day are added randomly to the spaces of the environment at a constant rate. These resources can represent food but can also represent resources in general including water, territory, and the access to those resources.

An organism’s amount of movement varies across the day following a circadian activity rhythm, which is modeled as a cosine wave with two parameters: amplitude, ranging from 0 to 1, and phase angle, ranging from 0° to 360° (Fig 1). The number of moves performed between times *t*_*0*_ and *t*_*1*_ is:

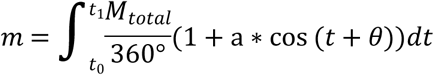

where *m* is the number of moves, *M*_*total*_ is the total number of moves per day = 20, *a* is the amplitude, and θ is the phase angle. Moves were calculated in 10 timesteps per day (*t*_*1*_ - *t*_*0*_ = 36°). We believe a sinusoidal activity rhythm is a reasonable approximation, and a more flexible activity pattern would in fact make temporal segregation easier to achieve. Though an animal’s activity profile varies according to its circadian rhythm parameters, the total number of moves is fixed at 20 moves per day; we hold total movement and energy expenditure constant so that the primary variable of interest is is the timing of movement. Each animal has the capacity to store energy; 2 units of energy is expended per day for movement and metabolism, and 0.5 energy is gained when the animal consumes a resource. Animals die if their energy reaches zero, or if they reach 12 days of age. Values for arena size, energy expenditure, resource value, and death age are somewhat arbitrary, and were chosen at reasonable numbers which limited computational load.

### Reproduction

Depending on the scenario, simulations are run either with one or two species in the environment. Throughout the day at a constant rate, pairs of animals belonging to the same species are randomly chosen to mate. The probability of an individual being chosen for mating is proportional its energy; animals which have acquired more energy thus tend to have better fitness. During each mating, one offspring is produced, to whom each parent donates 1/3 of its energy. In some scenarios, the circadian amplitude and phase of all animals remain fixed at the same values, while in other scenarios, there is variation in the circadian parameters, which are acted on by natural selection to cause evolution. In the case of variable parameters, the child’s amplitude is the mean of the parents’ ± 0.04 SD, and the child’s phase angle is the circular mean of the parents’ ± 0.04*360° SD. Matings occur at a frequency such that the population of animals would increase by 10% each day if no animals died. In our two-species simulation scenarios, when not testing competitive exclusion, we stabilized the population levels so that random population fluctuations would not cause one species to disappear by chance. For this, a target population ratio was set (usually 1:1), and the species with a population below target reproduces slightly more, and the species with a higher population reproduces slightly less, proportional to the difference in population.

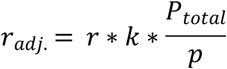

For each species, *r*_*adj.*_ is the adjusted population growth rate, which depends on the default growth rate *r* = 0.1, *k* the target fraction of the population which is composed of this species (*k* = 0.5 for a 1:1 ratio), *p* the population of the this species, and *P*_*total*_ the population of all species. The number of progeny in a timestep *t*_*0*_ to *t*_*1*_ is thus:

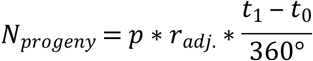

Calculations were performed in 10 timesteps per day.

### Assortative mating

To study speciation, the standard simulation parameters were modified to model assortative mating. We assume that mating must occur in a 9-hour (135°) time window which is fixed relative to the organism’s overall phase angle of entrainment, and the probability of mating between two individuals is proportional to the amount of overlap of their mating windows, ranging from 0 to 1. In other words, individuals which have a phase difference of nine hours or more do not mate, and the probability of mating increases linearly as the phase difference approaches zero. We define a “cross-group mating” metric to measure reproductive segregation in the population. To calculate this, we first define an overlap matrix of mating windows, i.e. the matrix where each cell *M*_*ij*_ is the mating-window overlap between individuals *i* and *j*. K-means clustering is used on the overlap matrix to define two clusters of individuals. The cross-group mating index is the mean of the overlap values between individuals from different groups; this is equivalent to the probability that a given mating is between different groups. Cross-group mating of 0 thus indicates complete reproductive isolation of two groups.

## Supplementary information captions

**S1 Dataset. Population sizes over time in competitive exclusion tests.** Dataset for Fig 2. Sheets of the Excel file are labeled A-C, and each sheet contains population levels of two species over 1200 days in 40 trials. (A) Organisms have amplitude = 0. Dataset for Fig 2A. (B) Organisms have amplitude = 1, and the two species have the same phase angle. Dataset for Fig 2B. (C) Organisms have amplitude = 1, and the species have a phase angle difference of 180°. Dataset for Fig 2C.

**S2 Dataset. Phase angles during character displacement.** Dataset for Fig 3. Mean peak-activity phases for two species over 400 days in 20 trials.

**S3 Dataset. Amplitude of a single species over time.** Dataset for Fig 4 A-C. Mean population amplitude for a single species over 8000 days. 20 trials were performed for each of three specialization values: 0.1, 0.4, and 0.75.

**S4 Dataset. Relationship between amplitude and phase variance.** Dataset for Fig 4D. Amplitudes (representing intraindividual niche expansion) were progressively stepped down from 0.99 to 0. For each amplitude increment of 0.01, 500 samples of the standard deviation of phase angle (representing interindividual niche expansion) were taken.

**S5 Dataset. Amplitude of two species.** Dataset for Fig 5 A-B.Sheets of the Excel file are labeled A-B, and each sheet contains the mean population amplitudes of two species over 25000 days in 20 trials. (A) Specialization was set to 0, and target population ratio between the two species was set at 1:1. One trial was chosen for Fig 5A. (B) Specialization = 0.1, population ratio 1:1. One trial was chosen for Fig 5B.

**S6 Dataset. Amplitude of two species with uneven populations.** Dataset for Fig 5 C-D. (A) Specialization was set to 0.1, and target population ratio between the two species was set at 2:1. One trial was chosen for Fig 5C. (B) Specialization = 0.1, population ratio 9:1. One trial was chosen for Fig 5D.

**S7 Dataset. Cross-group mating values during speciation.** Dataset for Fig 6. Cross-group mating (CGM) values over 1000 days in 40 trials.

**S8 Dataset. Evolution of amplitude versus speciation.** Dataset for Table 1. Amplitude and cross-group mating (CGM) values over 1000 days. Each sheet of the Excel file contains 20 trials in one combination of amplitude standard deviation and phase-angle standard deviation.

**S9 code. Python script for the simulation.**

